# Effect of cell size and tethering geometry on rotation rate, torque, and rotational bias of *E.coli* cells

**DOI:** 10.1101/2020.05.11.088310

**Authors:** Vidhu Soman, Kanishk Singh Malik, Sunil Nath, Ravikrishnan Elangovan

## Abstract

The bacterial flagellum has a rotary motor embedded in its membrane surrounded by stator-units. Torque is generated by electro-steric interactions between rotor and stator-units. Chemotactic signals entail the motor to switch its direction of rotation. However, other factors such as protonmotive force and torque are involved in switching. In this work, we used peritrichously flagellated E.coli that stochastically tethers on surfaces in random geometries and studied how the cell size and position of the rotational axis affect the output of the motor. We developed a Cell Tethering Analysis Program (CTAP) to measure the length of the cells, the axis of rotation, rotational frequency of tethered cells. A D/L ratio (diameter traced by the cell body to the length of the cell body) was used to quantify the location of the rotational axis wherein, a D/L ratio of 1 and 1.9 means the axis of rotation is at the center of the cell body and near the tip of the cell body respectively. We performed experiments in controlled conditions and quantified the effect of cell size and tethering geometry on the output of the flagellar motor. The estimated torque of the tethered cells was 951 pN.nm and 1390 pN.nm for a D/L ratio of ∼1.0 and ∼1.9 respectively. As the torque increased, the motors rotated exclusively in CCW direction. We conclude that quantifying the cell size and tethering geometry is significant to characterize the output of the flagellar motors in a tethered cell assay.

## Introduction

Motility system of flagellated bacteria has a rotary motor and stators inside the cell, a filament (propeller) which extends from the cell, and a hook which connects the filament and the rotary motor^1,2^. The filaments of peritrichously flagellated microorganisms such as *E*.*coli* and *Salmonella* bundle and propel the cell forward^1,3^. The motor’s stator-units have proton channels, and proton flux through the channels powers the rotor by converting the chemical energy of the ion motive force to mechanical work ^4,5^. The motor does not require ATP^6^ or other ions^7^ for rotation. Viewed from the filament to the cell body, the motor rotates in counter-clockwise (CCW) or clockwise (CW) direction. The filaments unbundle when one or more filaments switch from CCW to CW direction that re-orients the cell body to a new path. Chemotactic signals switch the motor rotation by changing the concentration of a cytosolic CheY protein^8,9^.

One of the first evidences for rotary motion of flagella is by demonstration of rotating tethered bacterial cells^10^. When a flagellum tethers to a glass surface, the cell body rotates, which can be visualised with brightfield microscopy. Flagellum was tethered using anti-flagellin antibody coated on glass surfaces^10^. This assay has been simplified by the generation of mutant *E. coli* strains that express FliC^sticky^ flagellin, which forms sticky filaments that spontaneously bind to glass surfaces ^11^ and made the tethered cell assay easier to set up.

Torque of bacterial flagellar motor has been estimated by the tethered cell assay. The bacterial flagellar motor (BFM) operates in the low Reynolds number regime wherein viscous forces dominate and inertial forces are negligible^12^. Hence, the torque generated by the BFM is balanced by the viscous drag experienced on the cell body. Tethered *E. coli* cells with maximum pmf rotate at slow speed (10-15 Hz) due to high load of the cell body^13^. Torque generated by BFM was estimated using drag force calculations under varying viscosity of the medium and by electrorotation of cell body. Tethered cells generate constant torque at speeds up to 10 Hz ^14^. The rotational speed is changed by varying the viscosity of the medium, and the estimated torque was 2700 pN.nm ^15^ for swimming cells and ∼880 pN.nm^16^ and 2200 pN.nm^17^ for tethered cells. And electrorotation experiments consolidated these torque estimates ^18–20^.

Switching of flagellar motor from CCW to CW direction is not only due to the chemotactic stimuli but also due to other factors such as proton motive force and load experienced by the motor^21,22^. In swimming cells, the rate of switching is much higher compared to the tethered cells where the flagellar motor experiences high load, and the switching is sensitive to high load experienced by the motor where the speed is less than 50 Hz^22^.

In the tethered cell assay, the immobilised cells have various sizes and tethering geometry. Length of the cells and location of the rotation axis change the drag experienced by the motor which affects the rotation rate and torque^13^. Hence, to estimate the torque and rotation rate, the cell size and position of rotational axis must be quantified. Some methods use equipment such as photomultiplier tubes^13,19^ and linear graded filters^18^ to detect the position and size of the cell body. Another method uses additional long exposure image acquisition to determine the location of rotational axis^23^. Thus, data processing of tethered cells usually requires additional equipment and cumbersome image analysis. Lack of automated data analysis for tethered cell assay makes it time consuming. The output of tethered cell assay has significant variations due to the random immobilisation and non-specific interaction with the surface (unbound flagellum interacting with the glass surface). Such non-specific interactions cause cells to wobble during the data acquisition which creates uncertainty in the quality of the data. So, these random fluctuations should be accounted during data analysis.

We performed the tethered cell assay of *E*.*coli* in controlled conditions to develop a robust and automated tethered cell analysis program, to quantify the size and tethering geometry the cells and to study their effect on the rotational rate, torque, and switching of the tethered cells. We have developed a cell tethering analysis program (CTAP) that measures the change in angular position of cell body, the CCW and CW rotational frequencies, the duration of rotation in both directions, the length of the cell, the centroid of the cell body, and the location of rotation axis. The program requires only brightfield movies of tethered cells, and no additional image processing. It thresholds the movies, removes noise, and measures the parameters mentioned above. With these parameters, we calculated the torque of the tethered cells using the method as described before^24^. We quantified the change in frequency of rotation (and torque) with the cell length and the location of rotation axis. Using centroid data, we eliminated the rotating cells that wobbled (noise). We detail our tethered cell assay program and discuss the variations in tethered cell analysis due to the cell length, the location of rotation axis, and how it affects the rotation rate, torque, and rotational bias of tethered cells.

## Results

The tethering of peritrichously flagellated *E. coli* on glass surface is a stochastic process which changes the axis of rotation of cells. The position of rotational axis of the cell body affects the frictional load experienced by the flagellar motors. To study the effect of cell size and tethering geometry of *E. coli* in pH 7 and at 30 °C, we developed an automated cell tethering analysis program (CTAP) to quantify the cell length, the location of rotational axis, the frequency of rotation. The program can be downloaded from the GitHub link provided in the supplementary section.

We performed the tethered cell assay in pH 7 motility buffer to avoid any variables that affect switching rate and rotational frequency. Thus, changes in rotational frequencies in our data are due to the cell size and tethering geometry. The CTAP previews the movie which can be used to identify a suitable cell for analysis. Program automatically thresholds and analyse all images.

### i) User experience and robustness of the program

The output of the program (Table 1) was validated by simulating a rectangular object rotating with fixed angle (Fig.S7 and S8). The simulation was made with custom Matlab scripts (see supplemental information). The program measured the length of the object (Fig.S7) and the distance between the axis of rotation and centroid (Fig.S8) without any deviation.

**Table 1.**
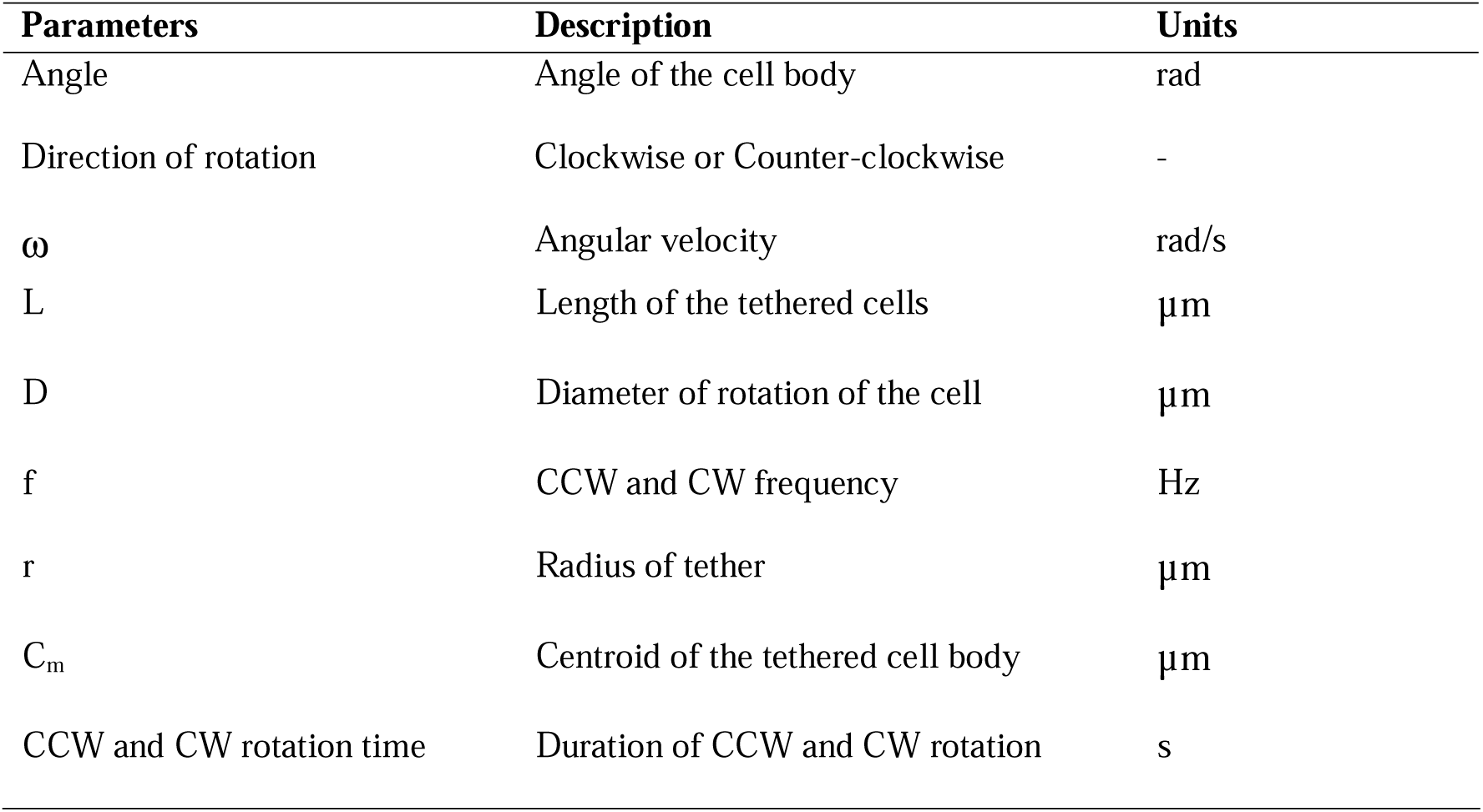
Parameters measured in the program with units.

| Parameters | Description | Units |
| --- | --- | --- |
| Angle | Angle of the cell body | rad |
| Direction of rotation | Clockwise or Counter-clockwise | - |
| $\omega$ | Angular velocity | rad/s |
| L | Length of the tethered cells | $\mu\text{m}$ |
| D | Diameter of rotation of the cell | $\mu\text{m}$ |
| f | CCW and CW frequency | Hz |
| r | Radius of tether | $\mu\text{m}$ |
| $C_m$ | Centroid of the tethered cell body | $\mu\text{m}$ |
| CCW and CW rotation time | Duration of CCW and CW rotation | s |

The program analysed 500 frames data in less than 20 seconds. We also analysed movies with 10,000 frames and the analysis duration was proportional (400 seconds). The graphical user interface and thresholding algorithm make the program easy to use, and it requires no programming knowledge. The real-time preview of the analysis shows the trace diameter which is useful to detect background noises (untethered cells) and wobbling of rotating cells. The program can be slowed down by adjusting the delay time (in the user input) for quality control of analysis.

We analysed 50 tethered cells. Some cells paused during rotation, and we used the output of the program to select the smoothly rotating tethered cells (26 cells). The centroid of tethered cells was used to identify cells with a uniform trajectory of rotation. Some cells wobbled (Fig.S5) which could not be detected under the microscope. When cells were tethered at the centre (C_R_ and C_M_ are closer or at the same point) the centroid data was clustered (Fig.S6C and D). Typically with long cells that were tethered at the end of the cell body, the centroid did not trace a circular trajectory (Fig.S6A and B). These cells were excluded from the final data compilation. We observed circular trajectory of centroid (stable rotation) when the axis of rotation was midway between the centre and the end of the cell body (Fig.S6E and F).

The program outputs the frequency of rotation per frame and cumulative change in angle (radians) for the assay duration (Fig.2A and 2B respectively). The change in angle reflects the switching of the motor (Fig.2B). The thresholding and noise removal algorithm used for image analysis removed the background noise in frequency data. Hence, we did not have to use any digital noise filters to smoothen the raw frequencies (Fig.2A).

**Figure 1.**
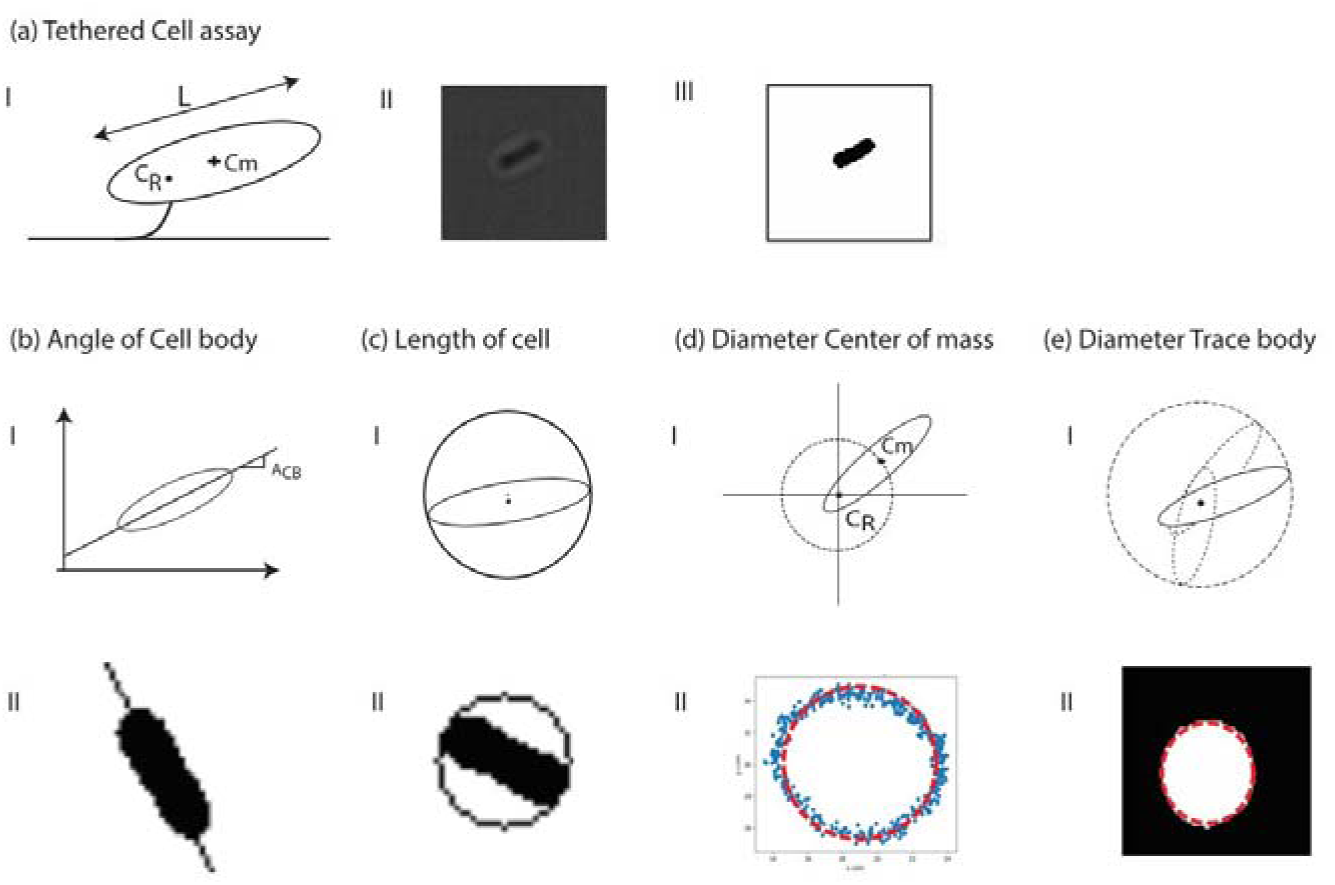
Parameters of tethered cells measured by the program. **A i)** Schematic of the tethered cell showing the parameters measured by the program. C_R_ is the axis of rotation, C_m_ is the centroid, and L is the length of the cell body. **A ii**) Bright field image of a tethered cell. **A iii)** Binarized image of a tethered cell. **B i)** Schematic of the angle of cell body with x-axis (detailed in methods**). B ii)** Linear regression analysis of a binarized frame of tethered cell. **C i)** Schematic of measurement of cell body length. **C ii)** Circumcircle around the is fitted to the cell body. The diameter of the circumcircle is used to calculate the length of the cell. **D i)** Schematic showing the trace of cell centroid around axis of rotation. **D ii)** Trajectory of centroid and circle regression fit. **E i)** Schematic showing the contour of the cell body rotation. **E ii)** Display of trace diameter calculated using edge fit.

**Figure 2.**
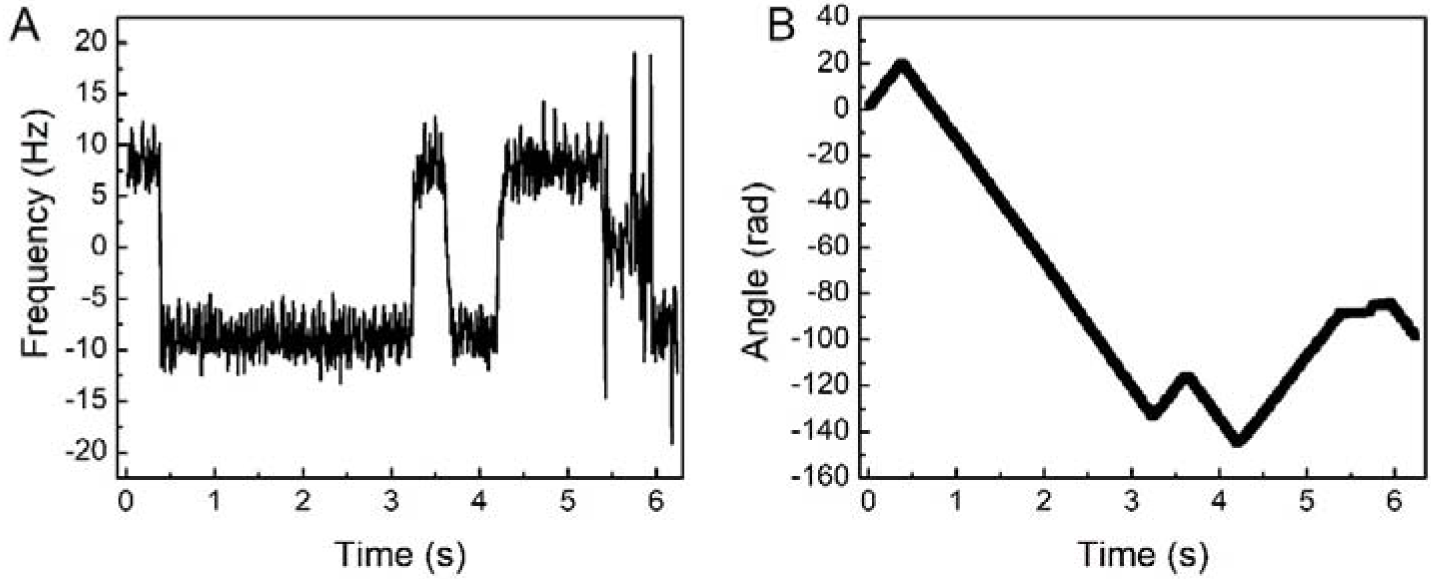
Representational data of frequency of rotation versus time (A) and cumulative change in angle (B) of tethered cell. Positive values represent CCW rotation and negative values represent CW rotation.

### ii) Measurement of size and tethering geometry of cells

The tethered cells were 2.5-4 µm long (Fig.3A), and the average length of the 26 cells was 3.2 ± 0.6 µm. The radius of tether in Figure 3B is the distance between the centroid and the axis of rotation. Radius of tether was used to find the length of long axis of two ellipsoids. Here, for shorter cells the tether was close to the centroid and for large cells, the tether was near the cell tip. We calculated the radius of tether using another method by measuring the trace diameter of cell body. Trace diameter of cell body is related to the radius of tether as D= L + 2r (Fig.S4A). Trace diameter ranges between L to 2L, depending on the position of axis of rotation (Fig.S4B). To define the location of rotational axis irrespective of the cell length, we used the ratio of the diameter of the circle the cell body traces (D) to the length the cell body (L). This dimensionless paramenter D/L can range from 1 to 2 as shown in Fig.3C and most cell are tethered close to the centroid of the cell. We observed a correlation of length of the cell with axis of rotation. Longer cells (> 3.2 µm) tethered near the end of the cell body (Fig.3B and 3D), while the shorter cells (< 3.2 µm) were tethered near the centroid of the cell body (Fig 3D). Cells longer than 4 µm were excluded from the data as their rotational trajectories were inconsistent.

**Figure 3.**
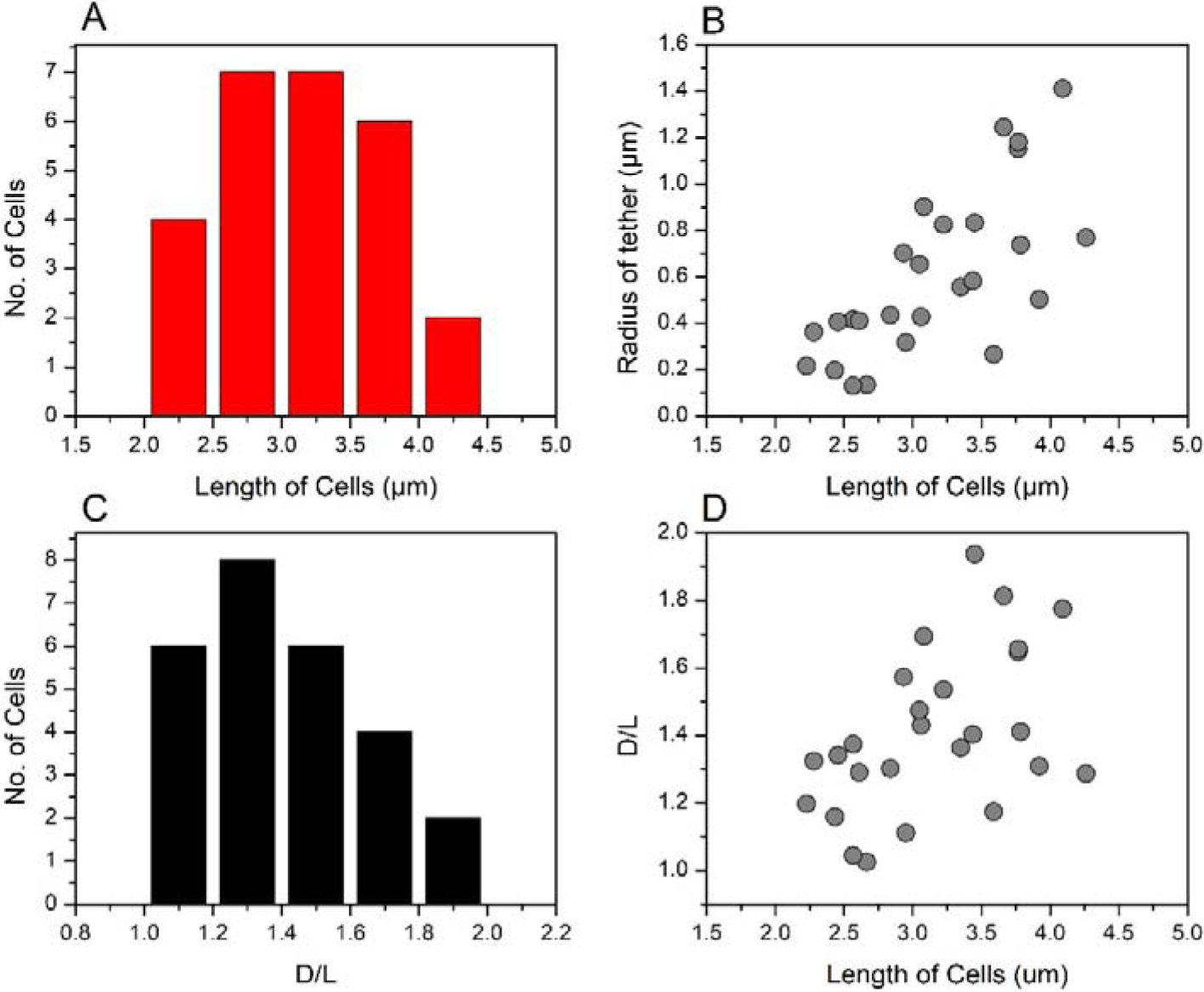
(A) Histogram of length of the cells analysed in this study. Length of tethered cells (L) varied from 2.2 µm to 4.2 µm. (B) Radius of tether versus the length of the cells. Radius of tether ranges from 0 to L/2 of cells. (C) Variation in D/L ratio of the cells analysed. D/L ratio varies from L to 2L. (D) Ratio of trace diameter,D, to cell length versus length of the cell. Cells tethered close to the centroid showed stable rotation. The data of 26 cells were included in this study. The average length of the cells was 3.2 ± 0.6 µm, average D/L ratio was 1.41 ± 0.24, and average radius of tether was 0.61 ± 0.35 (mean ± standard deviation).

### iii) Effect of cell size and tethering geometry on rotational frequency

The tethered cells rotated in both directions with equal magnitude (Fig.4D; R^2^ = 0.94). A maximum CCW and CW frequency of 13.95 Hz and 13.63 Hz respectively were measured from a data set of 26 cells (Fig.4). The frequency of rotation decreased with increase in cell length (Fig.4A. R^2^ = 0.70). Slow cells were 3-3.5 µm long, and shorter cells (2-2.6 µm) rotated at higher frequencies. The frequency of rotation decreased with increase in D/L ratio although the correlation was not linear (R^2^ = 0.3) (Fig.4B). As expected, the effect of the radius of the tether follows the same trend as the D/L ratio (Fig.4C). The frequency of rotation decreased with increase in r, as cells tether moves away from the centroid.

**Figure 4.**
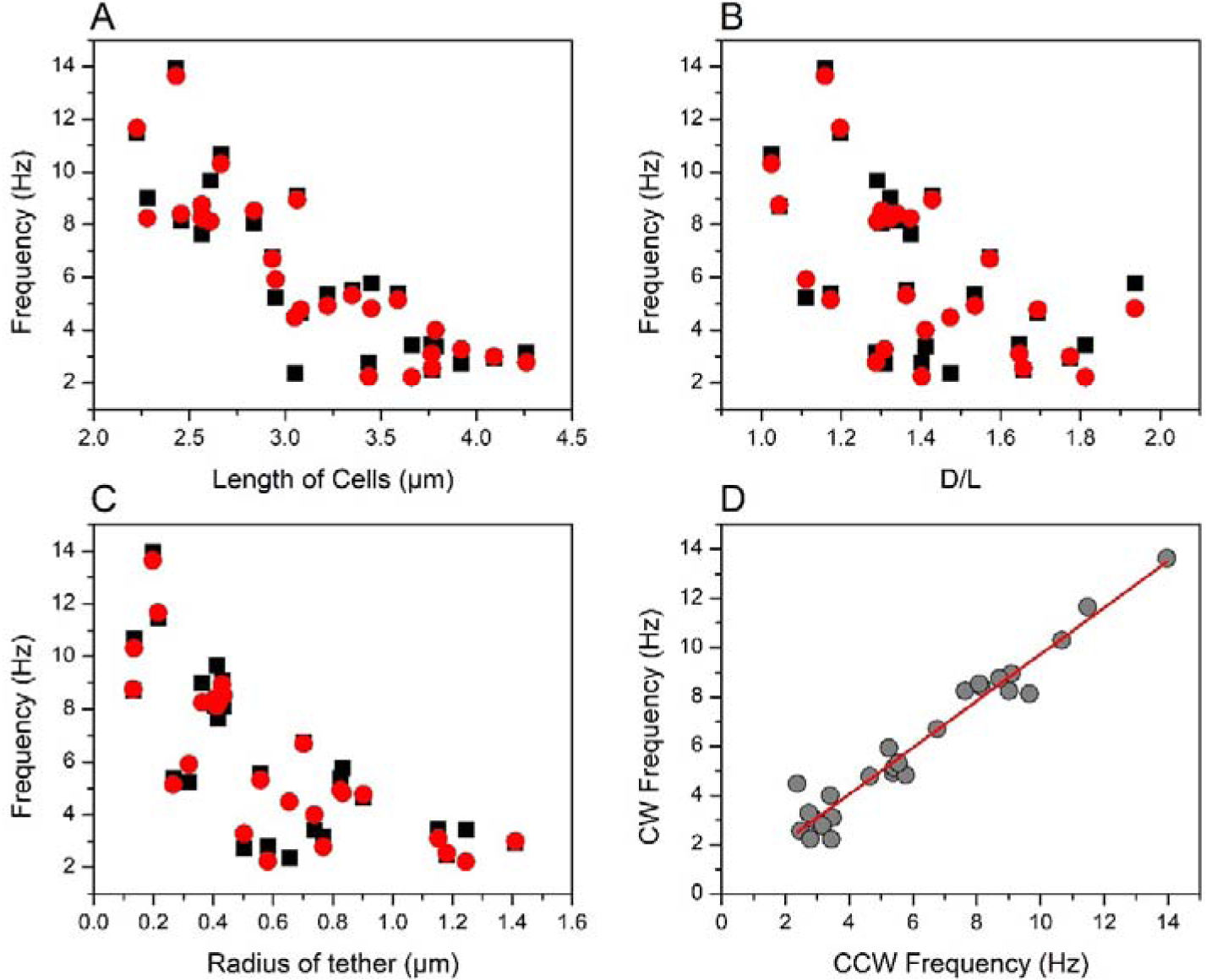
Effect of cell geometry on rotational frequency of tethered cells. (A) CCW and CW frequency is plotted as function of cell length. Red solid circles represent CW frequency and black solid squares represent CCW frequency. Longer cells rotated slower than shorter cells (R^2^ = 0.70). Number of cells included in the data was 26. The average CCW and CW frequencies were 1030.4 ± 268 pN.nm and 1008.1 ± 218 pN.nm respectively. (B) CCW and CW frequency is plotted as function of D/L ratio. (C) CCW and CW frequency is plotted as function of radius of tether. (D) CCW and CW frequencies of the tethered cells with linear fit (red line, R^2^ = 0.94). Rotational frequencies are identical in both the directions.

### iv) Effect of cell size and tethering geometry on torque

Torque did not show any linear correlation with the cell length. The average torque of the shorter cells (< 3.2 µm) was 955 pN.nm and of longer cells (> 3.2 µm) was 1118 pN.nm. Maximum CCW and CW torque generated by cells longer than 3.5 µm were 1483 pN.nm and 1517 pN.nm respectively (Fig.3B). A 4 µm long cell generated the maximum torque, which had a D/L ratio of 1.77. The motors generated more torque when the axis of rotation was near the end of the cell body (D/L > 1.6) in comparison to the cells with D/L ratio 1-1.6 (Fig.5B). The effect of the radius of the tether (*r*), which represents the distance between the axis of rotation and the centroid, on the estimated torque is similar to that of the effect of D/L ratio on the torque (Fig.5C). The torque speed curve of the motors analysed in this study was nearly constant (Fig.5D).

**Figure 5.**
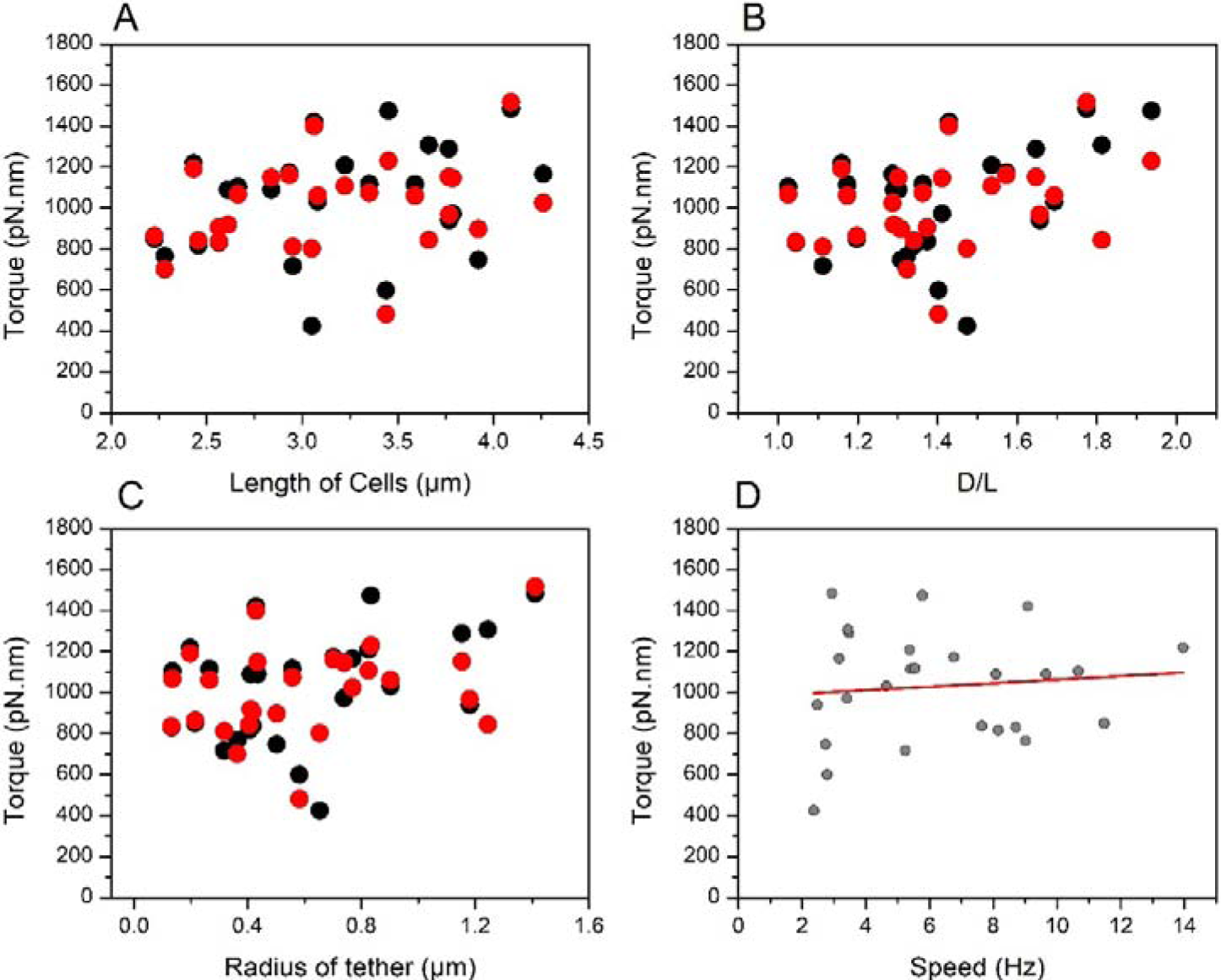
Effect of cell size and tethering geometry on torque of cells. (A) Change in torque as function of length of the cells. (B) Change in torque with D/L ratio, and radius of tether (C). The motor generated higher torque when the rotational axis was near the end of the cell body. Red solid circles represent CW torque and black solid circles represent CCW torque (D) Torque-speed curve of the motors analysed in this study. The change in CCW torque with the CCW frequency range 2-14 Hz. Red line is the linear fit of the data (R^2^= 0.1). Data of 26 cells included in this study. The average CCW and CW frequencies were 6.2 ± 3.1 Hz and 6.1 ± 3.1 Hz respectively.

### v) Effect of cell size and tethering geometry on rotational bias of the motor

The rotational bias had no linear correlation with the cell size, frequency of rotation (Fig.6A). Although longer cells rotated slowly, the cell length had no effect on the CCW bias (Fig. 6B). Cells that are tethered near the centroid are CW biased and the cells that are tethered near the tip are CCW biased (Fig.6C). When motors generated higher torque, the rotation was biased to CCW direction (Fig.6D).

**Figure 6.**
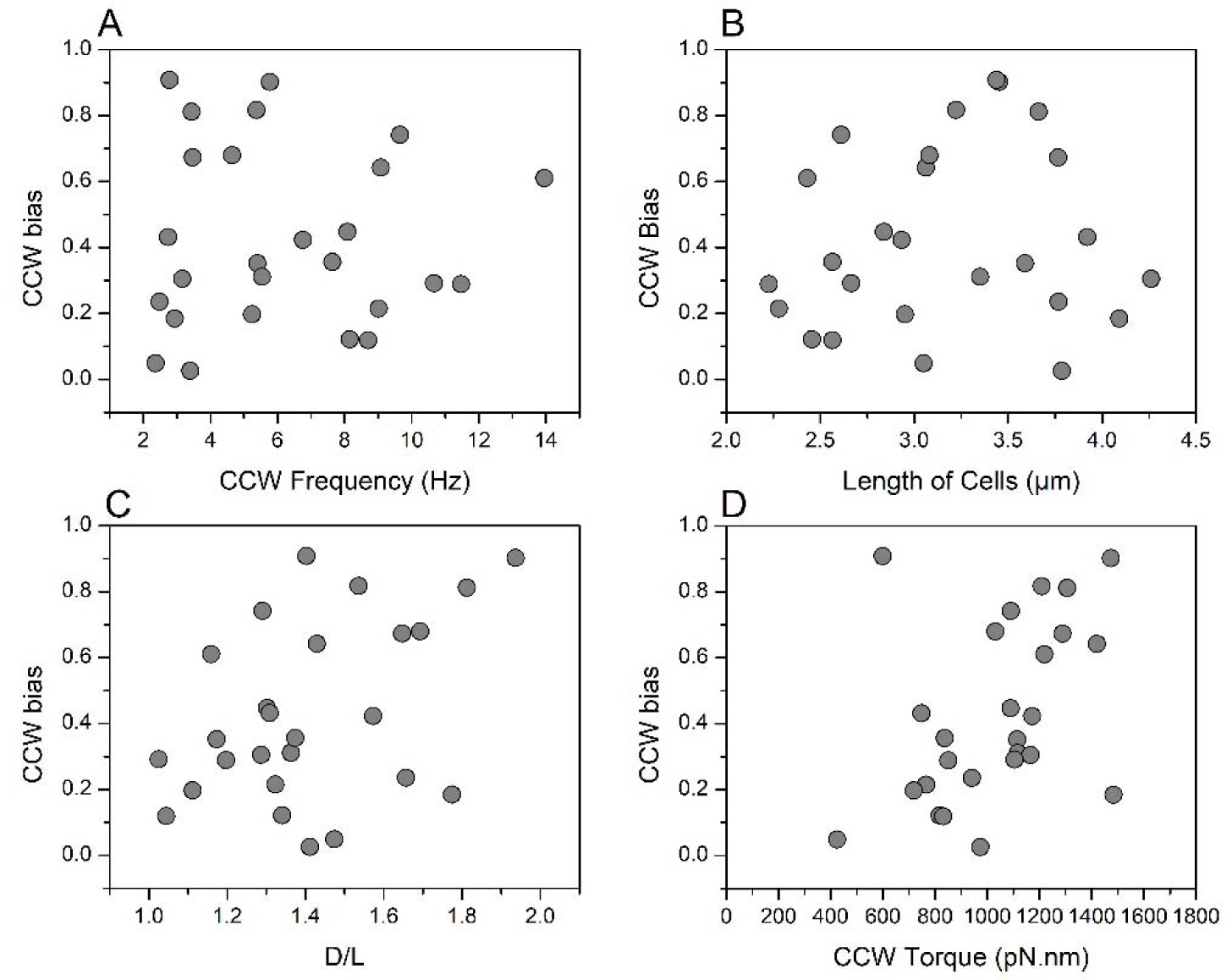
Effect of cell size and tethering geometry on CCW rotational bias of the motor. (A) CCW frequency versus CCW rotational bias of motor for the cells analysed. (B) Rotational bias and length of the cells. The length of the cells alone showed no correlation with the rotational bias. Motors generating higher torque rotates exclusively in CCW direction. (C) Change in CCW bias with D/L ratio. When the cells were tethered near the end of the cell body, the motor was CCW biased. (D) Change in CCW bias with CCW torque. Except torque, CCW bias showed no correlation with frequency, cell length, and D/L ratio.

## Discussion

We found that the axis of rotation and the cell body length of the tethered *E*.*coli* affect the output of the flagellar motor. We developed a Cell Tethering Analysis Program (CTAP) and measured the clockwise and counterclockwise rotational frequencies, length of the cell, radius of tether, and the position of rotational axis of cells. The program was developed in Python with graphical user interface and was completely automated. Unlike previous methods, the program analyses tethered cell movies from just brightfield movies and requires no additional equipment or data processing step. The program was used to identify wobbly and irregular rotating cells, and they were removed from the dataset.

Cell size and asymmetrical tethering cause fluctuations in rotation rate of tethered cells. Determining the axis of rotation has been an optical and image analysis challenge. The fluctuations have been detected by calculating the variance in angular position as the rotation rate varies at certain angular positions per revolution^25^. These variations are not due to the random fluctuations by the motor’s mechanism, or Brownian diffusion of cells^25^. Some studies eliminate the torque variations due to differences in length of cells by calculating relative torque^26^. In studies that use photomultiplier tubes to detect the position of the cell body, the transducer follows a point on the cell body^13^, close to the reference frame, or the centre of mass of the cell body^19^. However, these methods do not give information about the size and shape of the cell and is only useful for cells tethered near the end of the cell body. Also, such methods need linear graded filters in data analysis^18^. In another method, length of the cell was measured from microscopy images, and the rotation axis was measured taking long exposure images of cells. In long exposure, the cells appear as bright discs and the diameter of the disc is measured. Using this diameter and the length of the cells, the position of the rotation axis is determined ^23^. This method is like the D/L ratio method we used, but it needs additional steps in image acquisition. Our program uses only brightfield images to calculate all the mentioned tethered cell assay parameters. We analysed hundreds of movies and identified large percentage of cells that showed irregularities in rotation rate due to wobbling of cells which we consider as noise in the assay.

In tethered cell assay, rotation of the cell body is observed to study the flagellar motor. As it rotates in low Reynolds number regime, the cell body experiences viscous drag which equals the torque generated by the flagellar motor. The torque at which the cell body rotates is given as b⍰ω, where b is the frictional drag coefficient which depends on the size and shape of the cell, axis of rotation, and the distance between the cell and the surface, ⍰ is the viscosity of the solution, and ω is the angular velocity^27^. To estimate the size of the cell, we averaged the width of the cell body as 0.5 µm by manually measuring the width of cells using ImageJ program, which is close to the width of *E*.*coli*. The program measured the length of the cell (L) and calculated the location of rotation axis by fitting the centre of mass data (Fig 1d ii).

The torque-speed curve is a characteristic of a motor. For a given cell length, the change in rotation axis changes the viscous drag of the cell body which affects the torque generated by the motor. The motor senses the load and responds to changes in load^28^. The observed decrease in rotation rate of the motor with cell length is due to the increase in viscous drag. The drag is proportional to the cube of the length of cell body (see materials and methods). For instance, a 4 µm long cell tethered at the end will have a drag coefficient factor of 64. When it is tethered at its centre, the drag coefficient is 16. Thus, theoretically, for a given cell length, change in rotation axis from centre to the end of the cell increases the drag coefficient by 300 %.

Histograms in figure 3 show the total number of cells used in this study, the diversity of cell length, and location of rotation axis in our assay after eliminating cells with irregularities in rotation. As mentioned before, we analysed over 50 cells and the data of only 26 cells were used. That is, close to 50% of the cells were showed irregularities in rotation. This shows that, in a tethered cell assay, majority of the cells may not show stable rotation. Hence, analysis of the rotational trajectory is imperative to identify stable tethered cells. As the tethered cell assay is easy to set up, one can analyse hundreds of cells with the CTAP. However, the number of cells to be analysed depends on what needs to be studied. As the cells tethers stochastically on the glass surface, analysing more cells would not make any sense. We wanted to show the number of ‘bad cells’ in a tethered cell assay. It would not be reduced as we analyse more cells. The CTAP can be used to identify ‘good cells’ from the bad ones, which is why we targeted only 50 cells. Another aim of using the given size of the data set is to show how the load changes significantly in a tethered cell assay. That would not change even if the number of cells were increased. Majority of the cells are 3-3.5 µm long (Fig.3A) and are tethered close to the centre of the cell body (Fig.3C). We observed that longer cells have more propensity to tether near the tip of the cell body compared to shorter cells (Fig.3C). We quantified the distance between axis of rotation and centroid of cell body in two ways: Trace diameter (D) and radius of tether (r). We observed that the frequency of rotation of the cells changed with the length of the cell and the location of the axis of rotation. Irrespective of the rotation axis or cell length, the CCW and CW frequencies of a given motor were of same magnitude (Fig.4D) because of the symmetric mechanism of torque generation in both the directions^29^ (For a review see ^30^). When the length of the cells doubled, the frequency of rotation decreased ∼85 %. Due to high viscous drag of longer cells, the rotation rate decreases (Fig.4A). If the rotation axis of all the cells in the assay is same, then the rotation rate would change linearly with the cell length. However, cells tether stochastically on surface which changes the location of rotation axis. So, if the location of rotation axis is unknown, a given cell length would show variations in rotation rate. When the axis of rotation changes, the change in load affects the frequency of rotation. Tethered cells with rotation axis near the end of the cell body rotated slower as compared to the cells tethered close to the centroid of the cell body (Fig.4). Slow rotating cells generate higher torque, which is what we observed as well. Hence, the length of the cell and location of axis of rotation affect the rotation rate (and torque).To measure the torque of the tethered cells, it is imperative to calculate the frictional drag coefficient of the cell body which depends on the cell shape and tethering geometry. Except the distance between the tethering surface and the cell body, we measured all the parameters and calculated the torque.

We did not observe a corresponding increase in torque with change in length (Fig.5A) since rotation axis changes the viscous drag experienced by the cell body of given length. The torque of cells with D/L ratio <1.5 was lower than the torque of cells with D/L >1.5 as longer cells in our assay were tethered near the end of the cell body (Fig.5D). We constructed a torque-speed curve from our assay wherein the torque did not change linearly with speed (Fig.5D). For speed up to 14 Hz, the curve was almost flat. This is due to changes in load experienced by the motor because of variations in cell length and position of rotation axis of the cells. Variations in metabolic state among cells cause variations in rotational frequency and torque of the cells. We did not calculate relative torque (or frequency) because of the diverse cell length and rotational axes which affect the output of the motor. We acknowledge that the torque data in this study is based on an estimate by approximating the cell body as an ellipsoid. Hence, direct torque measurements on tethered cells with various rotational axes would give more insights into how the cells respond to changes in load.

The BFM stochastically switches the rotation direction. The rotational bias (CCW or CW bias) indicates the switching rate of the motor. Although a chemotaxis system causes the switching rate of the motor, torque generated by the motor is thought to affect switching, too^22^. In our assay, we calculated the CCW bias (see material and methods) to study the effect of change in viscous drag on switching of motor. It did not show any correlation with the cell length (Fig.6C) because of the change in axis of rotation. Also, the bias did not show any correlation to the frequency of rotation of motors (Fig.6A). The bias showed a correlation only to torque since torque is calculated from all these factors. We observed that the high torque, slow rotating motors were CCW biased (Fig.6B). Hence, to study the motor bias (or switching rate) of tethered cell system; cell length, rotation axis, and frequency of rotation must be considered.

## Conclusion

In tethered cell assay, the diverse cell size and random locations of rotation axis change the viscous drag experienced by the motor which would affect its output – rotational frequency, torque, and directionality of rotation. Without quantifying these factors, one would observe significant variations in the motor’s output even in control conditions. Quantifying these parameters usually requires additional equipment and data processing which makes the assay challenging. As our program quantifies the length of the cell and the rotation axis, we could identify the sources of variations due to the cell size and tether geometry. Load in tethered cell assay is usually assumed constant. However, change in rotation axis from the centroid of cell to the cell tip would significantly (100%) affect the viscous drag experienced by the bacterial flagellar motor. We observed that the length of the cells and location of rotation axis affect the frequency of rotation which must be factored in to study the output of tethered cells.

Our data shows that in a tethered cell assay, the stochastic tethering of the filament on the glass surface changes the load experienced by the motor. And, as the motors experience high viscous drag, they generate more torque to maintain a rotation rate. Hence, within a tethered cell assay, the load changes significantly. Direct measurement torque of the tethered cells would reveal how the motors’ torque change with load (with change in position of rotational axes). As the external load affects the incorporation of stators to the motor^31^, studying the stator dynamics in tethered cells with diverse axes of rotation would give insights into the same. Since we performed the assay in control conditions, the proton motive force was assumed to be constant for all the cells. Hence, the change in rotation rate is due to the increase in load which would mean that the load is a rate limiting factor in the rotary mechanism of BFM.

## Methods

### Strains

*E. coli KAF84* (a generous gift from Howard Berg’s lab) was used in this study. It is a wildtype for chemotaxis and has a plasmid with ampicillin resistance and a *fliC*^*sticky*^ gene, which produces flagellar filaments that stick readily to glass surfaces.

### Culturing and growth of strains

Tryptone broth (1% Bacto tryptone, 0.5% NaCl) was used to grow the strain. A loop full of culture from an agar plate was inoculated in 10 ml Tryptone broth supplemented with 100 µg/ml ampicillin. The culture was grown at 33 °C while being shaken at 200 rpm. The cells were harvested when the OD_600_ reached between 0.6 – 0.8. The culture was divided into ten 1 ml aliquots and washed twice with pH 7 motility buffer, which is 10 mM KPi, 0.01 mM EDTA, 70 mM NaCl. The washed cells were suspended in fresh motility buffer.

### Tethered cell assay

A flowcell was made by layering a square coverslip (22 mm × 22 mm) and rectangular coverslip (60 mm × 24 mm) between two strips of double sticky tape forming a channel of 24 × 8 mm and ∼ 20 µl volume. The cell suspension in pH 7 motility buffer was added to the slide and incubated for 15 minutes for tethering to the glass surface. The unbound cells were removed by washing it with fresh pH 7 motility buffer (∼50 µl). To prevent evaporation of the sample, a pool of fresh motility buffer was added on both the open ends of the slide. Around 3-6 motile tethered cells were observed in every field of view.

### Microscopy and Data acquisition

Olympus IX 71 inverted microscope equipped with a 1.4 NA 60X oil immersion objective, sCMOS camera (Andor Zyla), and halogen lamp for bright field illumination were used. The slide was mounted on a custom-built temperature controller maintained at 30 °C. A time series of bright field images were captured with 10 ms exposure time for 500 frames. The image area was 512×512 pixels and the size of the pixel was 0.122 µm. The movies were captured using ImageJ Micro Manager camera software.

### Data Analysis: Cell Tethering Analysis Program (CTAP)

#### Data handling, image processing, detection of cell body and its orientation

The folder with the time series of brightfield images in tiff format was browsed from the GUI of the program (Fig.S1 in the supporting material). After the input of the required parameters for (total number of frames, metadata availability, frame rate, and speed of real-time display), the program runs few frames for the user to select the cell. An area around the tethered cell of interest was cropped for image thresholding. The program read the images, stored the average intensity value of pixels, and the binary thresholding algorithm binarized the image (Fig.1a). After thresholding, the program applied noise removal filter and a median filter (removes salt and pepper noise) to remove noise. A dilate function joined broken areas in the region of interest which is due to difference in thresholding of the frames. The binarized pixel values were stored as an array of row and column co-ordinates. A linear regression fit on these pixel values was used to calculate the slope and intercept (Fig.1b). Using the slope, the angle between the regression line and the positive X-axis of each frame was calculated. Linear fit of the cell can be seen in real-time display.

### Calculation of change in angle of cell body, CCW and CW frequencies

The change in angle of cell body per frame is calculated by finding the difference in slope of consecutive frames (i^th^-(i-1) ^th^ frames) and stored in a vector A (using the NumPy tool). Then the function adds 180 to A (A_i_+ 180) and subtracts 180 (A_i_-180) and stored in vector B and C respectively. Another vector (D) is made whose i^th^ element is the minimum of the absolute values of the other vectors (A_i_,B_i_,C_i_). The directionality of rotation is stored in another vector, E, whose values are either +1 (CCW) or −1 (CW). The directionality of rotation is assigned as follows: If A_i_/D_i_ = 1 or B_i_/D_i_ = 1 or C_i_/D_i_ = 1, then it is +1, otherwise −1. The frame rate (obtained either from the metadata or from user input) and the change in angle was used to calculate the angular frequency (ω), and it is related to the rotational frequency as ω = 2πf rad/s, where f is frequency of rotation (s^−1^)

### Calculation of length of the cell

The program calculated the length of the cell by finding the diameter the circumcircle around the cell. It was a circle which completely covered the object with minimum area (Fig.1c). The diameter of the minimum enclosing circle was calculated for each frame and averaged to obtain the length of the cell (L). In cases where the standard deviation was high, the data was discarded.

### Calculation of radius of tether (r)

The distance between centroid of cell body and axis of rotation is called the radius of tether. The centroid of the cell body was calculated by averaging the X and Y coordinates of the pixels in binarized images using NumPy toolbox (a general-purpose array processing package). Plotting the centroid from each frame generates a circular trajectory (Fig.1d i). The program fits a circle to these points using *Least square method* to obtain the radius of tether *r* (Fig.1d ii). The centre of this circular fit was used as axis of rotation (C_R_). See table.S1 for the list of important functions used in the program.

### Calculation of trace diameter (D) of the cell body

The diameter of the trace of rotating cell body was calculated (trace diameter) by superimposing consecutive frames and calculated the contour. Contour was a curve which joined all the continuous points (along the boundary of the object) that had same pixel intensity. Contours were used in image analysis for shape dissolution, and object detection. The binarized images were inverted and superimposed to obtain a circular smudge of white pixels against a dark background, and the boundaries were used to find the diameter of the minimum enclosing circle (Fig 1e ii). Thus, the trace diameter changed from L to 2L depending on the tethering point of the cell (axis of rotation).

### Calculation of rotational bias

The program calculates the duration of rotation of the cell body in CW and CCW direction. The duration in each direction was summated to calculate the total time of rotation. The rotational bias was calculated as

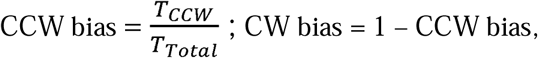

Where T_ccw_ is the duration of cell body rotation in CCW direction and T_Total_ is the total duration of rotation.

### Torque calculation of tethered cells

Torque produced by the tethered cell was calculated as described before^24^. Briefly, the rotating cell body was considered as two semi-ellipsoids around the rotation axis. The size of the ellipsoids depends on the position of the rotation axis (C_R_), length (L) and width of the cell. The program measured these parameters, and the frictional drag coefficient of large ellipsoid (*f*_*L*_) and small ellipsoid (*f*_*S*_) was calculated as follows:

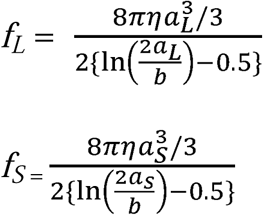

Where η = 9.6 × 10^−4^ Ns/m^2^, *a*_L_, *a*_S_, *b* were the viscosity of the motility media, major-axis length of the large semi-ellipsoid, the major-axis length of the small semi-ellipsoid and the minor-axis length of these two semi-ellipsoids (cell width) respectively. *a*_L_ and *a*_S_ were calculated by the following equations:

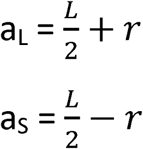

where *L* and *r* were cell length and radius of tether. The torque (*T*) was the product of frictional drag coefficient and angular velocity of the rotating cell and was calculated by *T* = ω(*f*_L_ + *f*_S_) pN.nm, where ω was the angular velocity in rad/s. The equations were solved with a custom script written in Matlab®.

## Authors contribution

VS & KSM did all the experiments and analysed the data. SN, VS & RE designed the study, reviewed the data, and wrote the paper. All authors reviewed the manuscript.

