## Supplemental Information for "Effect of cell size and tethering geometry on rotation rate, torque, and rotational bias of *E.coli* cells"

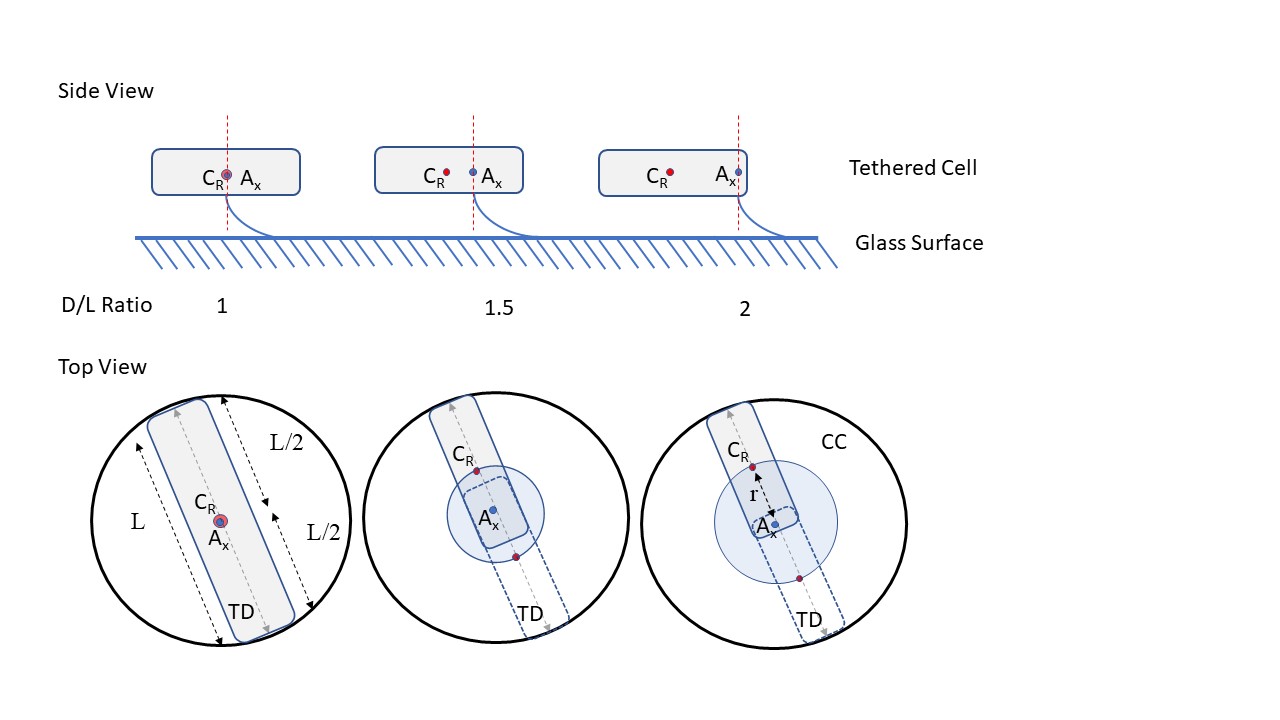

Figure S1. Illustration of top and side view of tethered cell showing D/L ratio (trace diameter to length of the cell body) with change in axis of rotation. CC is the circumcircle traced by the cell body (see materials and methods), TD is the trace diameter of the circumcircle. C_R_ is the centroid of the cell body, A_x_ is the axis of rotation (dashed red lines in the side view), L is the length of the cell, shaded blue circle is the trajectory of the centroid, and r is the radius of the tether. When the D/L ratio is 1, the axis of rotation is at the centroid of the cell body, and when the axis of rotation moves away from the centroid to the tip of the cell body, D/L ratio goes to 2 (top view).

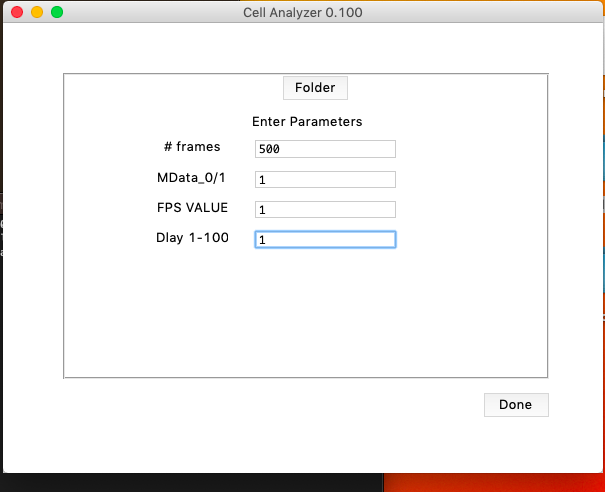

Figure S2. GUI of the program. The program prompts the user for number of frames (#frames), meta data (MData, 0 for no metadata and 1 if metadata is present), frame rate (FPS value. 1 for taking fps from the metadata), and delay time (speed of analysis. 1-100 is the number of frames analysed per unit time).

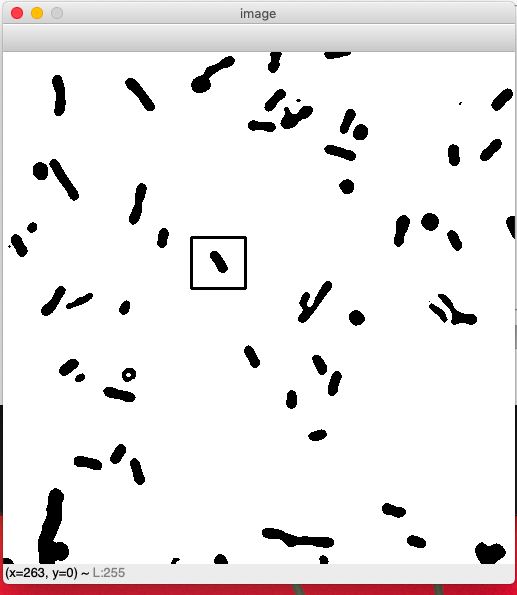

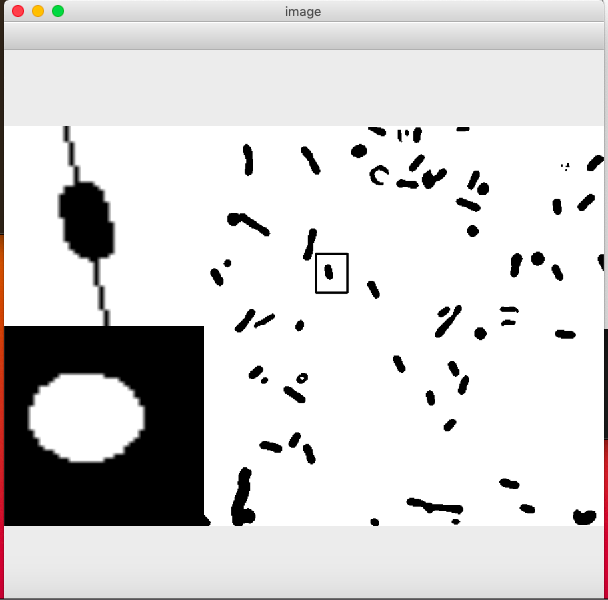

Figure S3. Cropping of tethered cell of interest from the binarized image (left). First, the program runs few frames of the movie. The cropped region is analysed, and the user can see the analysis in real time (right). The inset shows the linear fitting of the cell body (top left) and the trace diameter of the rotating cell (bottom left).

**Table S1. Important functions used in program**

| **Functions** | **Description** |
| --- | --- |
| *Threshold binary* | Thresholds the grayscale images |
| *cfilter2D* | Removes noise |
| *Median blur* | Removes salt and pepper noise |
| *Dilate* | Fix broken objects |
| *Imread* | Reads images |
| *Linear regression* | Uses simple linear regression equation. NumPy toolbox is used. |
| *Minimum enclosing circle* | Finds contours and calculates the trace diameter of the rotating cell body |
| *Least square optimize* | Calculates the radius of the circular trajectory of centroid. |

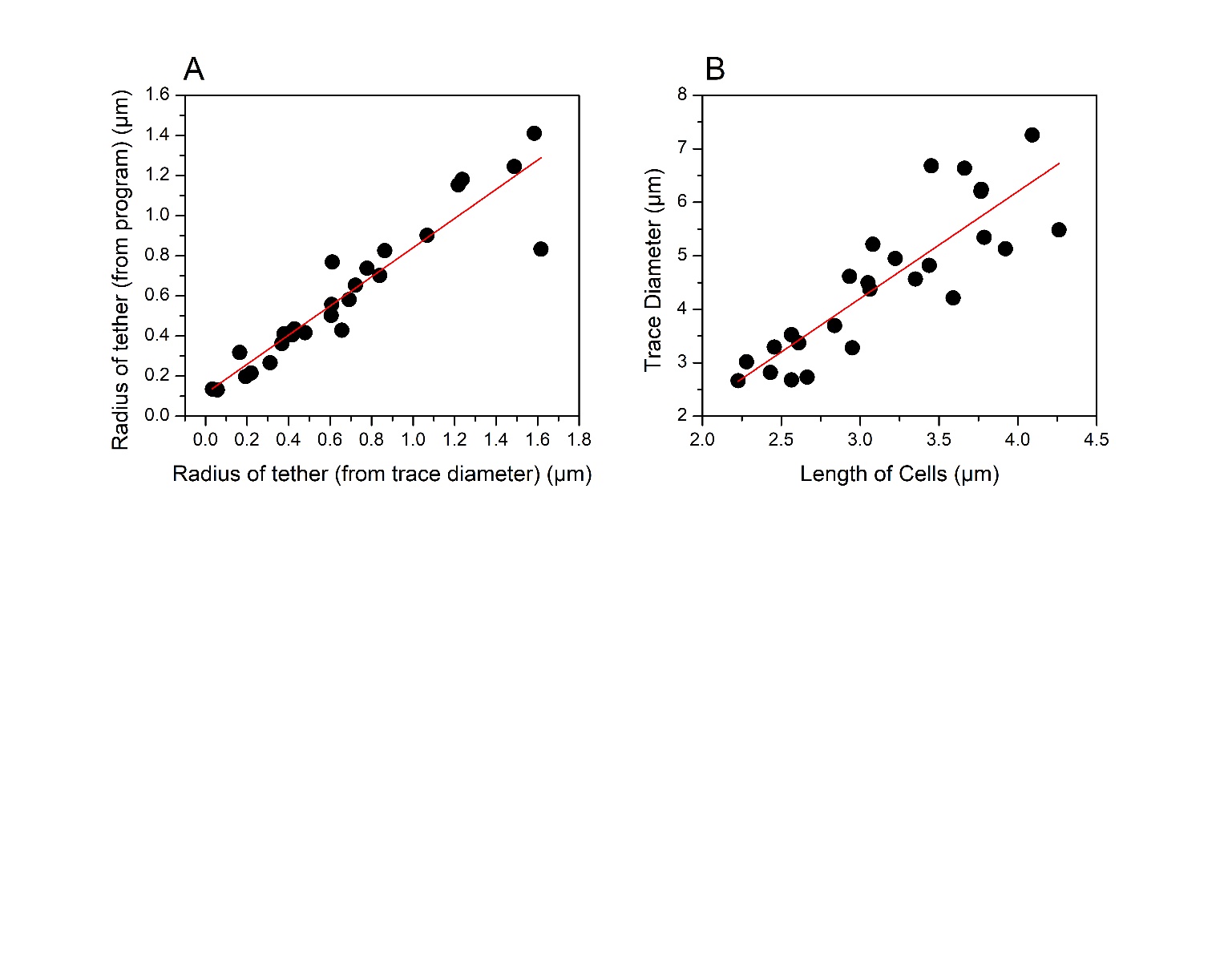

Figure S4: (A) Plot of radius of the tether measured in the program versus radius of the tether calculated from trace diameter. Red line is the linear fit of the data (R^2^ = 0.87). (B) Trace diameter versus the length of the cells analysed. The trace diameter becomes 2L for long cells as most of these cells tethered near the tip of the cell body. Red line is the linear fit of the data (R^2^ = 0.72).

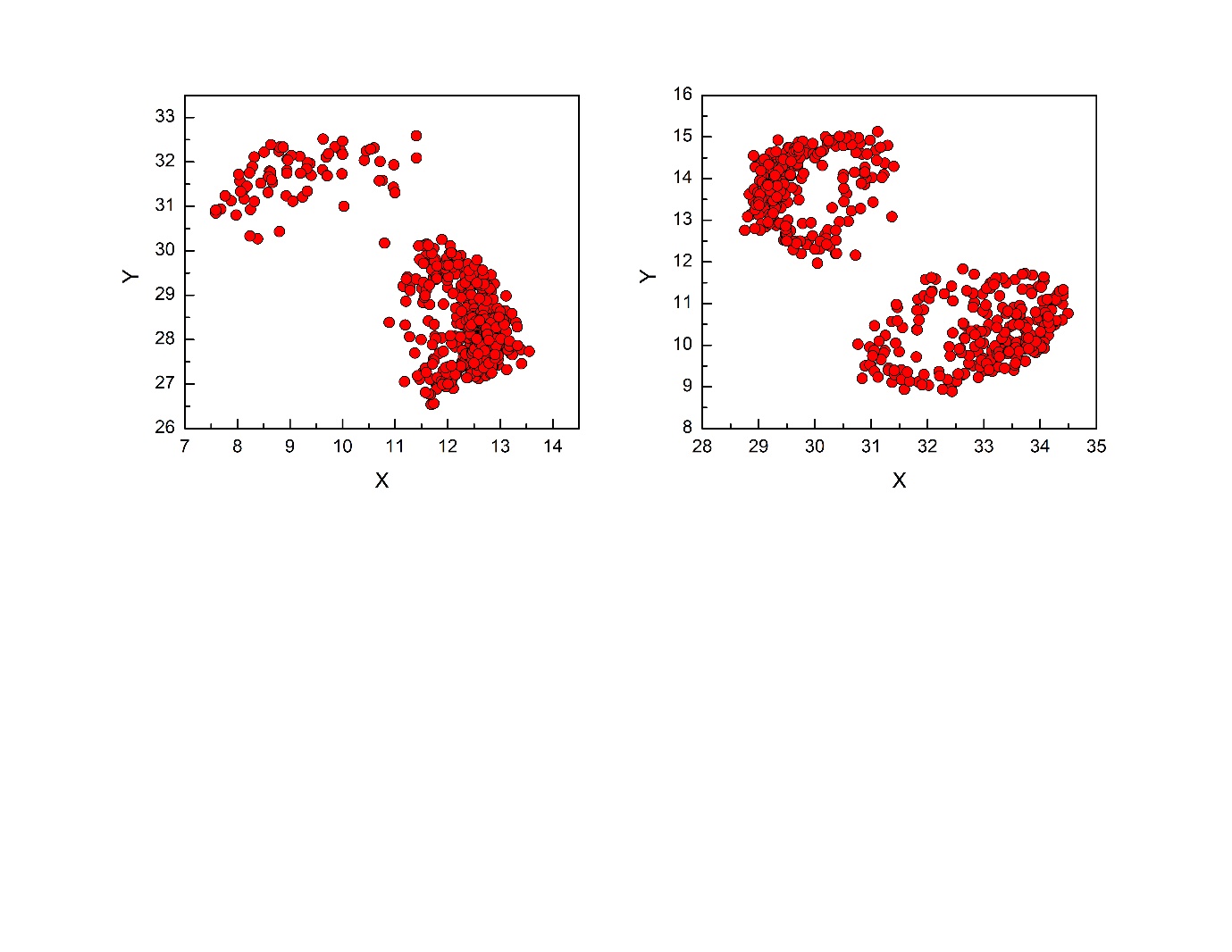

Figure S5. Centroid data of tethered cells that wobbled during rotation. These cells were removed from the data set as they showed irregularities in rotation.

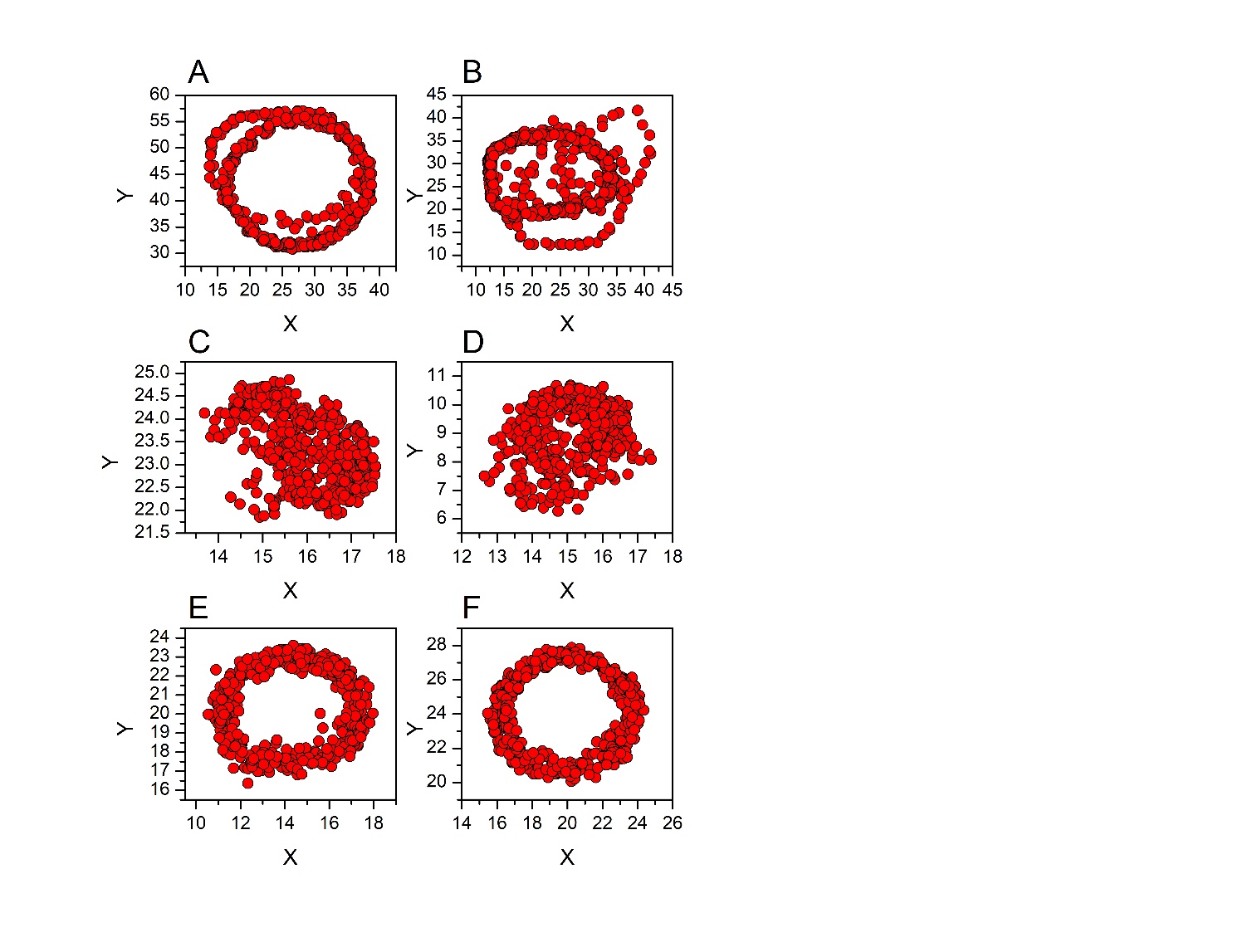

Figure S6: (A-B) Centre of mass data of cells tethered near the end of the cell body. The D/L ratios are 1.77 (A) and 1.81 (B). The length of the cells were 3.76 µm and 3.66 µm. When long cells tethered at the end, the centre of mass data would deviate from circular trajectory. (C-D) Centroid data of cells tethered at the centre of the cell body. The D/L ratios are 1.02 (C) and 1.0 (D). The centre of mass changed only 4-5 pixels in both the directions. (E-F) Centre of mass data of cells with intermediate D/L ratio (axis of rotation between end of the cell body and the centre). The D/L ratios are 1.33 (E) and 1.30 (F). Center of mass data has circular trajectory when the rotation axis is between the end and the centre of the cell body.

**Validation of data**

We made rectangular objects of various lengths and axes of rotation in Matlab to simulate tethered cells and to validate the size and tethering geometry using CTAP. A patch graphics object was drawn using the image processing toolbox of Matlab. The patch function drew the object with the specified *x* and *y* spatial co-ordinate system. For the background, a large white square object was drawn. Next, a black rectangular object was drawn inside this square to represent the cell body. This simulated the binarized images of the tethered cell assay. The co-ordinates of the vertices of the objects were specified and the patch function drew the object. The length of the rectangular object was changed by changing the *x*-coordinates of the vertices of a side. The length varied from 0.2 to 1 with 0.2 increments (spatial coordinates).

A rotate function in the Matlab was used to rotate the rectangular object. The function rotated the object by alpha degrees with a specified plane. The object was rotated around the z-axis, and the axis of rotation was changed from the centre of the object to its end. A fixed angle of rotation was specified and using a for-loop movies of 100 frames was made for analysis in the program.

These movies were analysed in the CTAP for the length of the rectangular object, radius, and D/L ratio. The output data from the program was linear to the input data and showed no variations (R^2^ = 0.99 for all the graphs in fig S7 and S8)

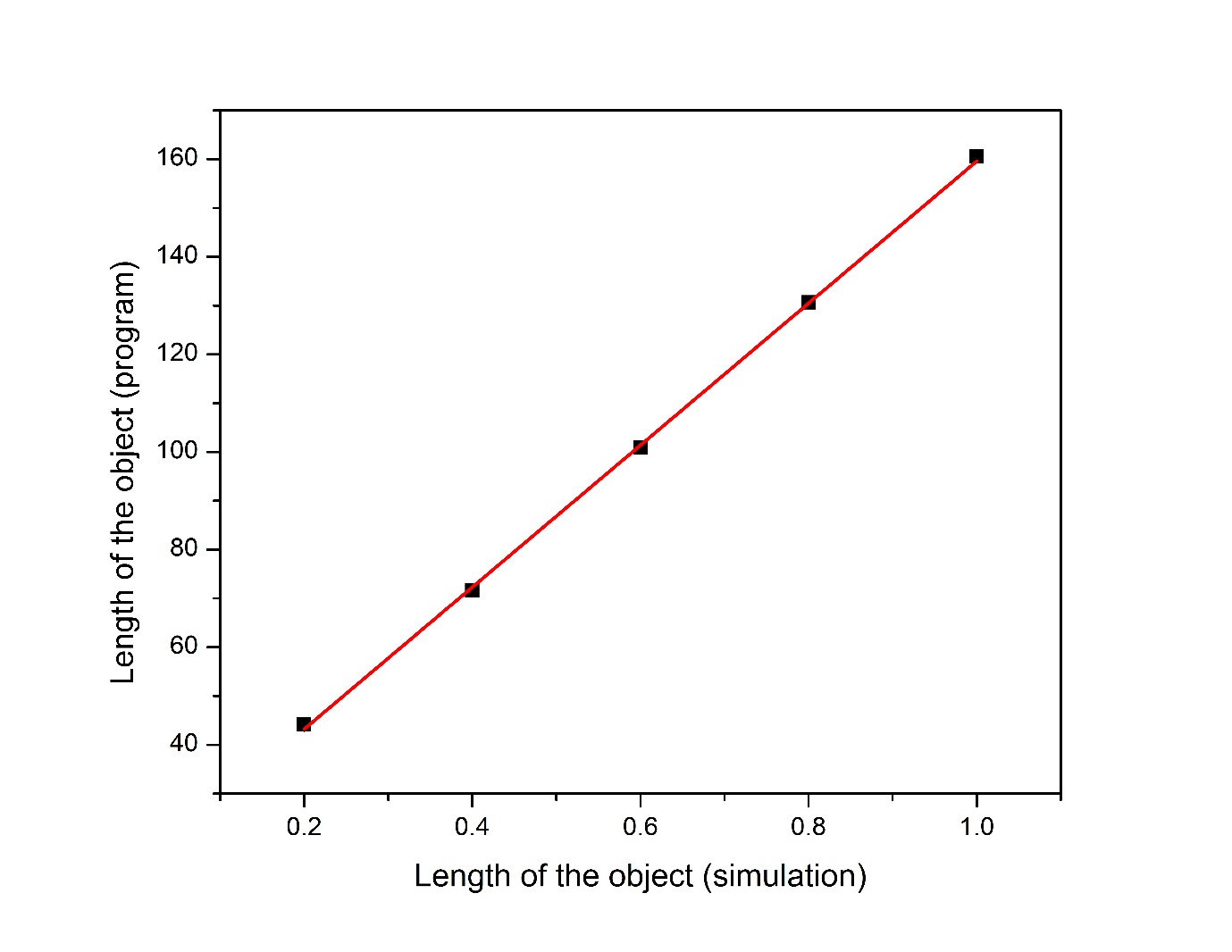

Figure S7. Length of the simulated object analysed by the program. Five movies with length ranging from 0.2 to 1 (arbitrary units) were made in Matlab. The length of object with different rotational axis were averaged and showed linearity (R^2^ = 0.99).

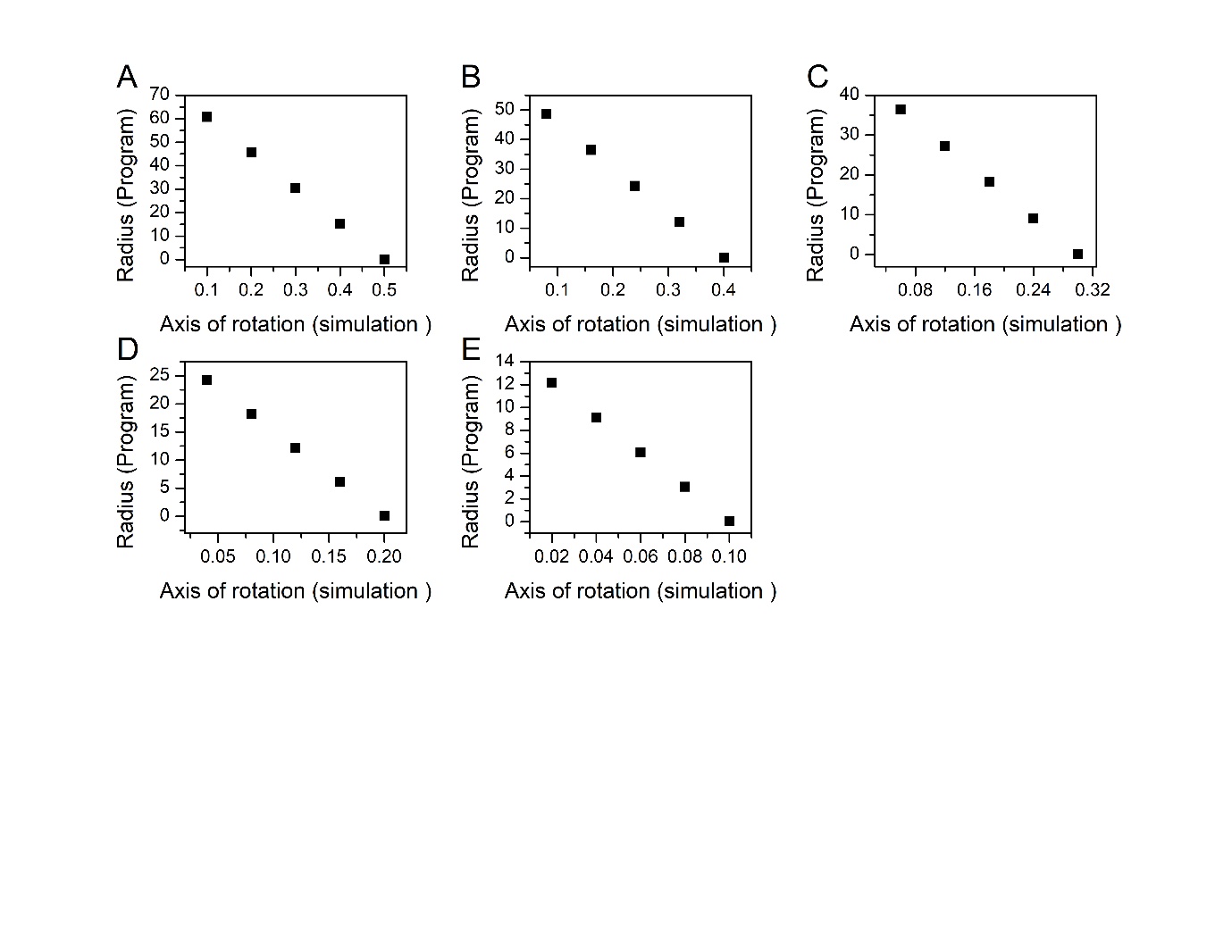

Figure S8. Radius and rotational axis of the object (A-E). Movies of the rectangular object with different axis of rotation were made in Matlab. A) Rectangular object with an arbitrary length of 1 B) 0.8, C) 0.6, D) 0.4, E) 0.2. The axis of rotation was specified in the simulation with increments shown in the graph. The radius (radius of the tether) was measured by the CTAP with no deviation from the known values.

The GitHub link to the tethering analysis program is <https://github.com/kanishk-ux/Tethered-cell-analysis>
